# Room to breathe: Nutrition and developmental oxygen modulate the crowding effect on size in *Drosophila melanogaster*

**DOI:** 10.64898/2026.07.02.736161

**Authors:** Cole M. Nicholls, Alexander W. Shingleton

**Affiliations:** Department of Biological Sciences, University of Illinois Chicago, IL 60607

**Keywords:** Developmental plasticity, nutritional plasticity, hypoxia, body size, larval crowding, *Drosophila*

## Abstract

In a wide variety of animals, developmental crowding results in adults with smaller bodies. The crowding effect on body size in *Drosophila melanogaster* is canonically attributed to heightened competition for nutrition. However, whether other consequences of crowding also contribute to its effect on size remains an open question. We tested the relative contributions of nutritional competition, oxygen availability, and larval-generated metabolites to the crowding effect on size. We found that while nutrition explains most of the variation in body size due to crowding, oxygen also contributes in a sex- and nutrition-dependent manner. We found no evidence that larval-generated chemicals affect body size. These data confirm a widely suspected but untested role of nutrition in producing the crowding effect on size in *D. melanogaster*, while revealing an unexpected role of oxygen, and raise the possibility that behavior may be a mediator of density-dependent plasticity.

**Research Highlights:** We found that both nutrition and oxygen mediate the crowding effect on size in *Drosophila melanogaster*.

## Introduction

Across a wide variety of animals, population density experienced during early life is known to alter an organism’s morphology, physiology, and life history (Wilbur 1980, Smallegange 2022). For example, the armyworm *Spodoptera exempta* develops a more melanized cuticle at high densities as a defense against infection (Reeson et al. 1998, Wilson et al. 2001), while tiger salamander larvae living at high density develop into cannibals with broader heads and larger teeth than typical larvae (Rose and Armentrout 1976, Collins and Cheek 1983). While some density-dependent effects are species-specific, others are widespread. In particular, in many animals, individuals developing in more crowded conditions grow into smaller adults (Matte et al. 2025). However, because this crowding effect on size (hereafter, CES) has been documented largely in wild populations where manipulating developmental conditions is challenging (Albon et al. 1983, Grant and Imre 2005, Beukema and Dekker 2015), the mechanisms underlying CES in many cases remain poorly understood. Addressing this gap therefore requires a genetically tractable model in which density-dependent development can be studied experimentally, such as *Drosophila melanogaster*.

For over a hundred years, researchers have been studying the effects of developmental crowding in *D. melanogaster* (Pearl and Parker 1922). Larval crowding has been shown to affect a variety of adult traits, from developmental time (Sang 1949, Bakker 1962) and lifespan (Miller and Thomas 1958) to thermotolerance (Sørensen & Loeschcke 2001) and fecundity (Lints & Lints 1971). *D. melanogaster* larvae also exhibit CES: larvae reared in more crowded conditions develop into smaller adults (Alpatov 1929, 1932, Sang 1949, 1950, Bakker 1962). Canonically, this effect has been attributed to heightened competition for nutrition (Sang 1949, 1950, Bakker 1962), and recent work demonstrated that supplemental yeast increases body size in larvae reared in crowded conditions (Klepsatel et al. 2018). However, the role of nutrition in mediating the CES in *D. melanogaster* has not been fully tested. Further, whether nutritional competition is the sole explanation for CES, or whether other consequences of crowding contribute, remains unknown.

One potential but less obvious regulator of CES in *D. melanogaster* is developmental hypoxia. In almost all animals, hypoxia is a potent suppressor of body size (Harrison et al. 2015). *D. melanogaster* larvae may be exposed to hypoxia because, while foraging, they must balance the competing demands of access to oxygen and access to food. Larvae respire through two pairs of spiracles, one anterior and one posterior, that must remain exposed to air to maintain gas exchange (Manning and Krasnow 1993). Which spiracles are available for gas exchange depends on how a larva is feeding. Larval feeding can be grouped into three distinct behaviors (Kim et al. 2017): surfacing, in which both pairs of spiracles remain exposed to air; digging, in which their heads are buried in the medium but their posterior spiracles remain exposed to air; and diving, in which the larva descends fully below the surface with both pairs of spiracles retracted and respiration stops. Larval feeding behavior is therefore modulated by both the need for oxygen and the perceived availability of nutrients: blocking a larva’s posterior spiracles reduces digging behavior, while olfactory cues below the surface promote diving (Kim et al. 2017). If crowding depletes the nutrients near the surface of the medium, larvae may dive more frequently, increasing their exposure to developmental hypoxia and ultimately reducing adult body size.

Another factor that may contribute to CES in *D. melanogaster* is the accumulation of larval-generated metabolites in the medium. Larvae deposit metabolic waste products, cuticular hydrocarbons, and other chemicals whose concentration may increase in proportion to larval crowding (Bakker 1962, Henry et al. 2018). These compounds may reduce growth directly, by acting as a toxin, or indirectly, by acting as a cue indicating impending nutritional competition, analogous to how insects use environmental cues such as photoperiod to anticipate future changes in temperature (Saunders 2014). While recent work has shown that supplementing the larval diet with ammonia failed to replicate the effects of crowding (Henry et al. 2020), whether the full complement of larval-generated metabolites may affect CES remains unclear.

Here, we hypothesize that nutrition, oxygen, and larval metabolites mediate CES in *D. melanogaster*. To test the involvement of nutrition and oxygen, we manipulated levels of nutrition, oxygen, and larval density in a three-way factorial experiment. If nutritional competition drives CES, diet and density should interact such that high nutrition relieves CES; if hypoxia drives CES, then high oxygen should relieve it. Our results partially supported these predictions. We found that high dietary yeast relieves CES at all oxygen levels, whereas high oxygen only relieves CES in males reared on low dietary yeast. To test the involvement of larval-generated metabolites, we added small amounts of liquefied food from crowded vials to low-density vials. We found no evidence that this treatment affected body size, suggesting that larval-generated metabolites do not play a role in CES.

## Materials and methods

### Drosophila *stocks*

All flies were from the strain *Samarkand* (Bloomington Drosophila Stock Center #4270), and were maintained in 12”x12”x12” mesh chambers on standard yeast-sugar-carrageenan medium (**Supplementary Table S1**) at 25°C and 21% O_2_ unless otherwise stated.

### Egg collection and larval transfer

Flies were allowed to oviposit for 6-7 hours on a 10cm ⌀ Petri dish containing standard yeast-sugar medium spread with yeast paste. Eggs were left for 48 hours after the end of the lay, after which they were individually transferred to experimental vials as described below.

### Measurement of pupal case size

In all experiments, larvae were left to develop until their wings were sclerotized (P12-13). Pupae were then removed, sexed, and stored in 70% EtOH at RT until measurement. To measure pupal case size, the dorsal aspect of each pupa was imaged using a Leica DM6B equipped with a camera, and the area of the case was measured using ImageJ.

### Nutrition, oxygen, and density treatments

In our first experiment, we tested the effects of oxygen level and nutrition on CES using a fully crossed factorial design for all possible combinations of:

- Low nutrition (1:16 protein:carbohydrate diet) or high nutrition (2:1 protein:carbohydrate diet) (**Supplementary Table S1**);
- Hypoxia (15% O_2_), normoxia (21% O_2_), or hyperoxia (40% O_2_);
- Low density (50 larvae per vial), medium density (125 larvae per vial), or high density (200 larvae per vial).

We generated three vials per treatment combination.

In our second experiment, we tested the effects of hyperoxia on CES in males reared on low nutrition again using a fully factorial design but with a more focused set of conditions: low nutrition; normoxia or hyperoxia; and low or medium density. We generated eight vials per treatment combination. In both experiments, larvae were transferred to vials randomly with respect to environmental condition. In the first experiment vials were reared in three sequential blocks to account for potential temporal variation in rearing conditions. Low and high oxygen conditions were generated by rearing flies in a Coy Basic Glove Box with oxygen regulated by a Sable Systems Roxy-1 Universal Regulator.

### Larval-generated metabolites treatment

To test whether components of the larval-conditioned medium contribute to body size, we first generated conditioned medium by transferring 250 larvae to vials containing 7mL standard yeast-sugar medium. Six days after transfer, after all surviving larvae had pupated, we transferred either 0.1g or 0.2g of this liquefied media to low nutrition vials. Control vials contained only low nutrition food. Three vials were generated per condition. We then transferred 50 larvae to each of these vials. After pupation, pupae were removed and measured as described above.

### Statistical analysis

All data and R scripts used for analysis are available from Dryad (link). Pupal case size was log transformed prior to analysis and all data were confirmed to satisfy assumptions of normality and homoscedasticity.

To test the effects of oxygen, dietary yeast, larval density, and sex on pupal case size in the initial experiment, we fit the model: *P_ijklmn_* = *O_i_ • N_j_ • D_k_ • S_l_* + *V_m_* + *ε_ijklmn_* (model 1), where *P* is pupal case size, *O* is oxygen level (categorical fixed effect), *N* is nutrition (categorical fixed effect), *D* is density (categorical fixed effect), *S* is sex (categorical fixed effect), *V* is vial (random intercept), and the subscripts refer to the level of each parameter. We also tested whether including rearing block as a random factor improved model fit using a likelihood ratio test; as it did not, block was excluded from the final model.

To test whether hyperoxia modulates the crowding effect specifically in males, we conducted a follow-up experiment and combined this data with data from the first experiment that matched the conditions of the follow-up experiment. We then fit the model: *P_ijklm_* = *O_i_ • D_j_ • S_k_* + *E_l_* + *V_m_* + *ε_ijklmn_* (model 2), Experiment (*E*, categorical fixed effect) was initially included as a fixed effect to account for between-experiment variation; however, it was not significant, and was excluded from the final model.

To test whether components of larval-conditioned medium contribute to pupal case size, we fit the model: *P_ij_* = *T_i_* + *S_j_* + *V_k_* + *ε_ijkl_* (model 3), where *T* is treatment (categorical fixed effect), *S* is sex (categorical fixed effect), and *V* is vial (random intercept).

For all models, terms that were not statistically significant were removed, the model was refit, and the ANOVA was rerun on the simplified model. All analyses were conducted in R (version 4.6.0). Linear mixed models were fit using the *lme4* package (Bates et al. 2015), with degrees of freedom and *P*-values for fixed effects estimated using Satterthwaite’s method via the *lmerTest* package (Kuznetsova et al. 2017). The significance of fixed effects in all models was evaluated using Type III ANOVA with Satterthwaite’s method, and all ANOVA tests were conducted using the *stats* package (R Core Team 2013). *Post hoc* contrasts of estimated marginal means were conducted using the *emmeans* package and corrected for multiple comparisons (Lenth, 2024). Generative AI (OpenAI’s ChatGPT-5.5 and Anthropic’s Claude Opus 4.8) was used to assist the writing of R code for statistical analyses and in the copyediting of this manuscript.

## Results

Our data supported the hypothesis that density effects on body size are mediated by nutrition. Specifically, we found a significant interaction between the effects of nutrition and density (**Table 1**) such that CES was stronger under low-yeast conditions (**Fig. 1**). Our data did not support the simple hypothesis that density effects on body size are mediated by oxygen, as we observed neither a significant interaction between the effects of oxygen and density on body size nor a main effect of oxygen on body size (**Supplementary Table S2**). However, a *post hoc* analysis revealed that in males reared on low dietary yeast, high oxygen significantly decreased CES relative to low oxygen (**Supplementary Fig. S1**). This suggests that CES may be mediated by oxygen, but only under moderate crowding on low dietary yeast, and in a sex-specific manner.

**Figure 1.**
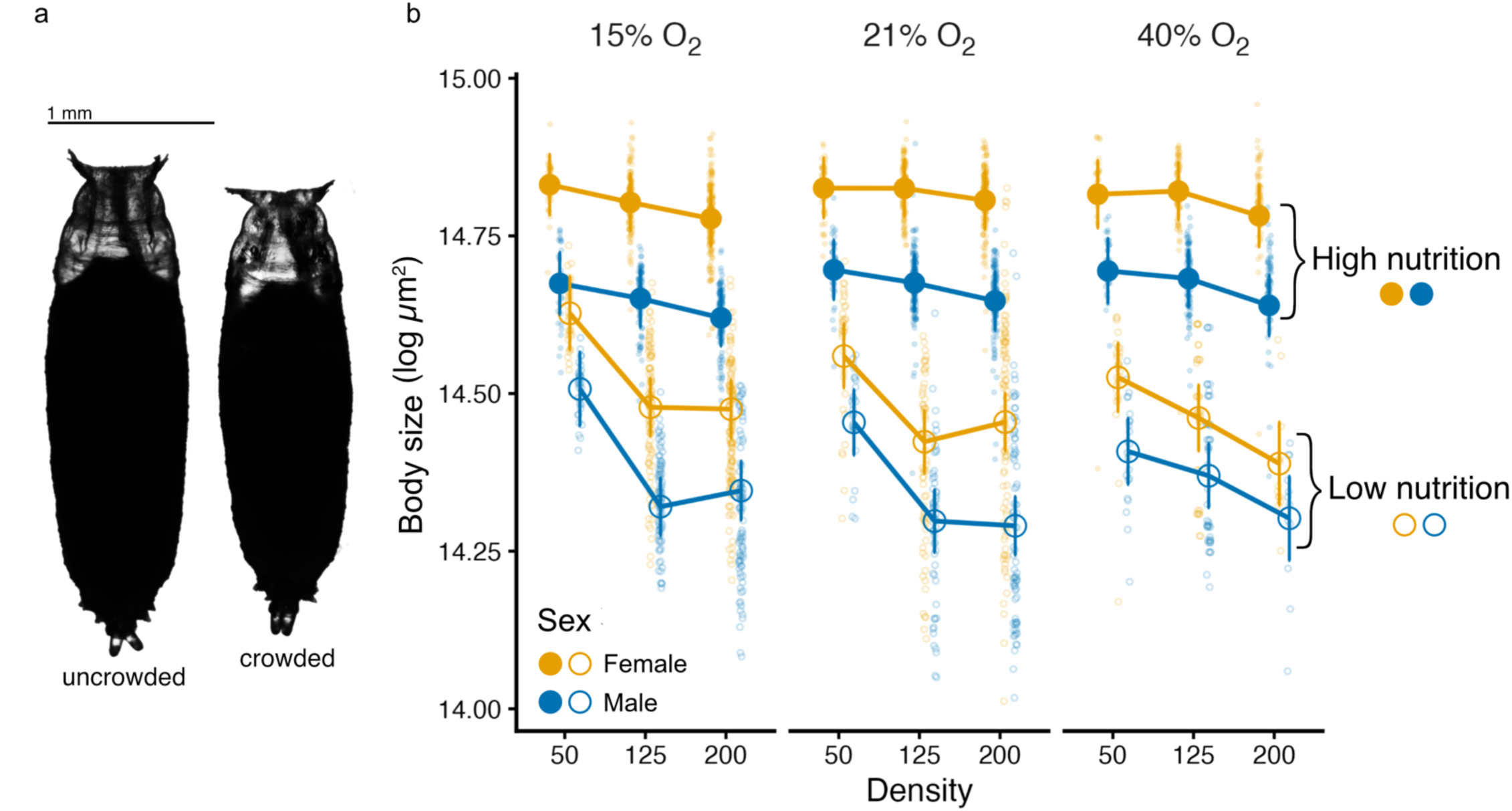
The effect of density, oxygen, nutrition and sex on body size in *D. melanogaster.* (a) Representative pictures of the pupal cases of male larvae reared in normoxia and low dietary yeast in uncrowded (50 larvae per vial, left) and crowded (200 larvae per vial, right) conditions. (b) Effect of larval density on body (pupal case) size across nutrition, oxygen, and sex conditions in the initial experiment. Large points show estimated marginal means and 95% confidence intervals (Model 1). Small points show individual flies. Closed points are high nutrition diet and open points are low nutrition diet. Female is orange, male is blue. Different panels reflect different oxygen conditions.

**Table 1.** - Results of ANOVA on the effect of density (50, 125, 200 larvae per vial), oxygen (15%, 21%, 40% O_2_), nutrition (low and high nutrition diets) and sex on body size (model 1), reduced to only significant terms.

| **Term<sup>†</sup>** | **SS** | **df<sub>Num</sub>** | **df<sub>Den</sub><sup>‡</sup>** | **F-ratio** | **P-value** |
| --- | --- | --- | --- | --- | --- |
| oxygen | 0.0077 | 2 | 37.3 | 0.67 | 0.516 |
| nutrition | 4.5855 | 1 | 37.6 | 805.92 | **< .001** |
| density | 0.2588 | 2 | 37.1 | 22.74 | **< .001** |
| sex | 10.5520 | 1 | 2,703.4 | 1,854.56 | **< .001** |
| oxygen:nutrition | 0.0393 | 2 | 37.3 | 3.46 | **0.042** |
| nutrition:density | 0.0995 | 2 | 37.1 | 8.74 | **< .001** |
| oxygen:sex | 0.0620 | 2 | 2,695.4 | 5.45 | **0.004** |
| nutrition:sex | 0.0465 | 1 | 2,699.0 | 8.17 | **0.004** |

<sup>†</sup> Type III sum-of-squares ANOVA. Model was stripped to include only significant terms. Original parameters included oxygen level, nutrition level, larval density, sex, and all interaction terms.
<sup>‡</sup> Denominator degrees of freedom (df) estimated using Satterthwaite's method.

In order to test this modified hypothesis, we conducted a targeted follow-up experiment in which we reared eight additional vials of larvae on low dietary yeast and tested the effects of two levels of crowding and oxygen (densities 50 and 125, and norm- and hyperoxia). An analysis of these data, combined with the data from our initial experiment, confirmed the three-way interaction between oxygen, density and sex (**Table 2**). We found that when larvae are reared on low dietary yeast, hyperoxia reduces the effect of crowding on male body size by over 50% relative to normoxia, while this effect is not observed in females (**Fig. 2**).

**Figure 2.**
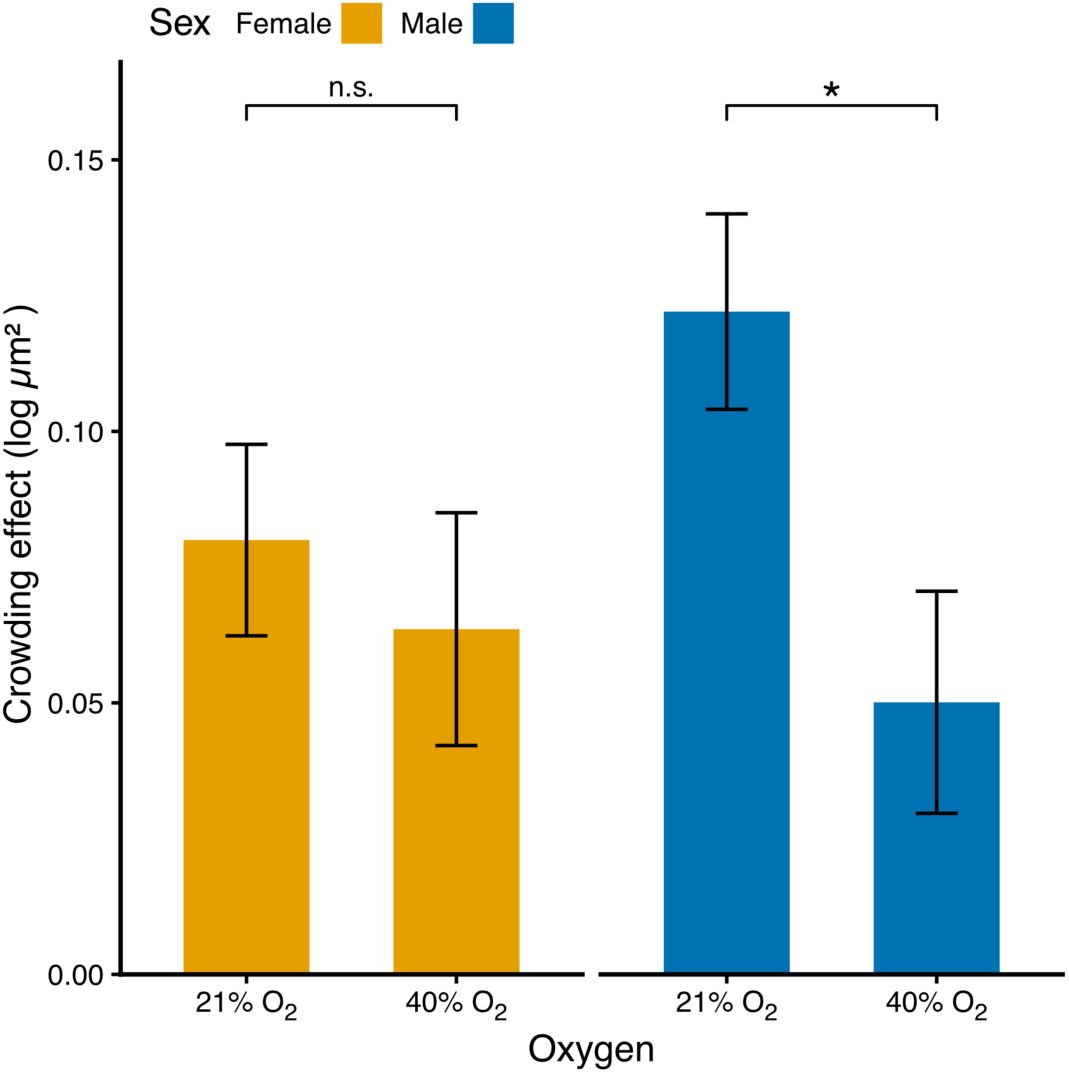
Hyperoxia reduces the effect of CES in males but not females reared on low nutrition diet. Crowding effect is difference in body size (estimated marginal mean, Model 2) for larvae reared at 50 versus 125 larvae per vial. Bars are SEs. n.s *P*>0.05, * *P*<0.05.

**Table 2.** - Results of ANOVA on the effect of density (50 v. 125 larvae per vial), oxygen (21% v. 40%) and sex on body size in larvae reared on low nutrition diet (model 2), reduced to only significant terms.

| **Term<sup>†</sup>** | **SS** | **df<sub>Num</sub>** | **df<sub>Den</sub><sup>‡</sup>** | **F-ratio** | **P-value** |
| --- | --- | --- | --- | --- | --- |
| oxygen | 0.151 | 1 | 40.8 | 14.191 | **<.001** |
| density | 0.464 | 1 | 40.8 | 43.530 | **<.001** |
| sex | 3.899 | 1 | 1,243.2 | 366.081 | **<.001** |
| oxygen:density | 0.037 | 1 | 40.8 | 3.431 | 0.071 |
| oxygen:sex | 0.130 | 1 | 1,243.2 | 12.209 | **<.001** |
| density:sex | 0.013 | 1 | 1,243.2 | 1.179 | 0.278 |
| oxygen:density:sex | 0.047 | 1 | 1,243.2 | 4.406 | **0.036** |

<sup>†</sup> Type III sum-of-squares ANOVA. Model was stripped to include only significant terms. Original parameters included oxygen level, larval density, sex, and all interaction terms.
<sup>‡</sup> Denominator degrees of freedom (df) estimated using Satterthwaite's method.

Finally, to test whether the crowding effect on size is mediated by chemicals generated by other larvae, we exposed larvae to food supplemented with larval-conditioned medium. We found that the addition of larval-conditioned medium had no significant effect on pupal size (F_2,6.1_ = 2.34, p = 0.18).

## Discussion

The crowding effect on size in *D. melanogaster* has been attributed to competition for nutrition, but this hypothesis has not been fully tested. Further, whether other consequences of crowding contribute to CES remains underexplored. In this study, we investigated the relative contributions of nutrition, oxygen, and larval-generated metabolites to CES in *D. melanogaster*. We found that nutritional competition is the primary driver of CES. Additionally, we found that oxygen plays a role in CES, although only in males reared on low nutrition. However, we found no evidence that larval-generated metabolites contribute to CES.

Previous work has consistently shown that crowded *D. melanogaster* larvae develop into smaller adults (Alpatov 1929, 1932, Sang 1949, 1950, Bakker 1962), and a recent study demonstrated that supplementing the larval diet with yeast relieves CES (Klepsatel et al. 2018). However, to establish nutritional competition as the mechanism underlying CES requires explicitly testing for an interaction between the effects of nutrition and larval density on body size; to our knowledge, no study has demonstrated such an interaction. We found that larvae reared on low nutrition experience much more severe CES than larvae reared on high nutrition, supporting the hypothesis that nutritional competition is a key mediator of CES.

Surprisingly, however, we found that in addition to nutrition, oxygen plays a role in mediating CES in males reared on low nutrition. One natural interpretation of this finding is that supplemental oxygen relieves competition for oxygen, in the same way that supplemental food relieves competition for nutrition. However, this explanation seems unlikely. Larvae were reared in vials with porous plugs through which gases can move freely, so any oxygen consumed by larvae should be rapidly replenished by oxygen from the surrounding atmosphere, eliminating competition. Why, then, do high oxygen levels relieve CES?

One possibility is that hyperoxia relieves CES in nutritionally restricted larvae by enabling them to access more of the food below the surface of the medium. To access these nutrients, larvae must perform dives during which they cannot respire (Kim et al. 2017). Supplemental oxygen may mitigate the costs of diving in two ways. Before a dive, higher oxygen concentration in the trachea may allow for longer or deeper dives; alternatively, after a dive, faster oxygenation of tissues due to higher atmospheric oxygen concentration might reduce the duration of hypoxic stress. Thus, hyperoxia may relieve the CES indirectly, by allowing larvae access to subsurface food, directly, by reducing the total hypoxic stress induced by diving, or both. Effectively, hyperoxia may expand the larval feeding band, the volume of food actually available to developing larvae (Venkitachalam and Joshi, 2026).

A second possibility is that hyperoxia relieves CES by increasing the activity of growth signaling pathways. Both moderate hypoxia and nutritional restriction reduce growth through shared signaling pathways (Kapali et al. 2022), so supplemental oxygen might partially restore pathway activity when nutrition is limiting. However, while hypoxia can reduce growth signaling pathway activity (Reiling and Hafen 2004), there is no evidence that hyperoxia can increase it. Consistent with this, developmental hyperoxia does not increase adult body size in *D. melanogaster* under standard laboratory conditions (Harrison and Haddad 2011, this study).

A third possible explanation is that the effect of hyperoxia on CES may be due not to phenotypic plasticity, but to viability selection. Viability selection occurs when environmental stressors cause non-random juvenile mortality with respect to a particular trait (Grafen 1988, Chippindale et al. 1993). If low nutrition increases variation in body size independent of oxygen level, and smaller males are more likely than larger males to die when reared on low nutrition and in hyperoxia, this would increase mean body size without affecting growth in any individual larva. To test whether viability selection could account for hyperoxia’s effect on CES, we included vial-level survival rate as a covariate in our combined model. When we conducted an ANOVA on this model, the three-way interaction between oxygen, density, and sex remained significant, suggesting that the buffering effect of hyperoxia on CES in males is not an artifact of viability selection (**Supplementary Table S3**).

None of the above hypotheses explain why the effect of hyperoxia should be specific to males. This result is surprising considering that female body size is more nutritionally plastic than male body size (Shingleton et al. 2017, Millington et al. 2021, McDonald et al. 2021). Given that moderate hypoxia and nutrition affect size through shared growth pathways, one might expect that hyperoxia would relieve CES more in females. Future work on the role of oxygen in CES should investigate the physiological or behavioral basis of this sex-specificity.

Another unexpected finding was that hypoxia did not elevate CES under low nutrition (**Table 1**). Indeed, hypoxia did not reduce body size at all, in contrast with myriad studies showing that rearing *D. melanogaster* larvae in 10% and 5% O_2_ reduces body size (Peck and Maddrell 2005, Harrison and Haddad 2011, Callier et al. 2013, Kapali et al. 2022). This indicates that our hypoxia treatment, rearing larvae from mid-L2 in 15% O_2_, may not have been sufficiently severe to affect size.

We found no evidence that the addition of larval-conditioned medium to vials of larvae at low density reduces body size. This result is consistent with recent work showing that the application of metabolic wastes like ammonia and urea do not replicate crowding effects (Henry et al. 2020), and suggests that chemicals generated by larvae are not a major driver of CES. However, our conditioned medium likely contained residual nutrients as well as larval-generated chemicals, which could have opposing effects on body size, and may have obscured a metabolite effect. It is also possible that the amount of conditioned medium applied did not contain enough larval-generated metabolites to induce an effect on size. Therefore, we cannot rule out the possibility that larval-generated chemicals modulate CES.

This study underlines the complexity of phenotypic plasticity in two ways. First, while plasticity is often modeled as a function of a single environmental variable (Woltereck 1909, Hudak and Dybdahl 2023), our findings show that body size in *D. melanogaster* depends on interactions among nutrition, oxygen, larval density, and sex. Our study therefore adds to a growing literature that treats trait values as a consequence of non-additive, context-dependent interactions among multiple environmental variables, a phenomenon known as multivariate or multidimensional phenotypic plasticity (Westneat et al. 2019, Nielsen and Papaj 2022, Hudak and Dybdahl 2023). Second, if hyperoxia relieves CES by promoting diving, this suggests that the environment may affect an organism’s morphology not just by directly altering its physiology, but also by altering its behavioral repertoire (Nielsen and Papaj 2022). A challenge for future studies of developmental plasticity, then, is to design hypotheses and experiments that capture both its multidimensionality and the possibility that it is shaped by behavior.

## Acknowledgements

We thank the members of the Shingleton Laboratory for their advice on technical matters and feedback on the manuscript; specifically, thanks to Elizabeth Agolli for the catchy title. Stocks obtained from the Bloomington Drosophila Stock Center (NIH P40OD018537) were used in this study.

## Data Availability Statement

The data and R scripts that support the findings of this study are openly available in the Dryad Digital Repository (DOI:XXXXXX)

---

**Supplementary Figure S1.**
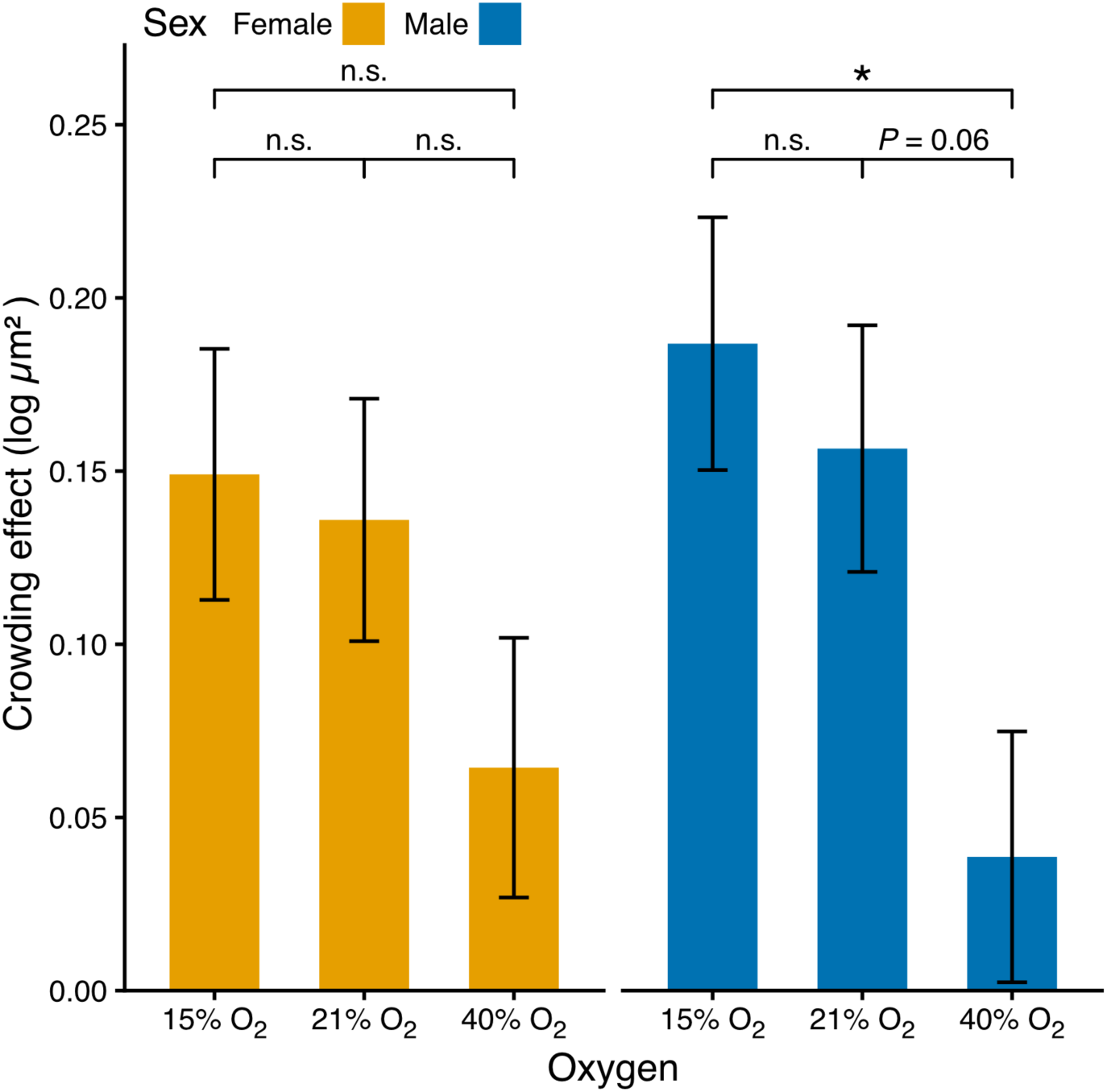
Hyperoxia reduces CES relative to hypoxia in males reared on low nutrition diet. Crowding effect is difference in body size (estimated marginal mean, Model 1) for larvae reared at 50 versus 125 larvae per vial. Bars are SEs. n.s *P*>0.05, * *P*<0.05.

**Supplementary Table S1.** Recipes for low and high-protein food used in the factorial experiments.

| P:C ratio<sup>†</sup> | Yeast:sucrose solution ratio<sup>‡</sup> | Yeast solution (ml)<sup>§</sup> | Sucrose solution (ml)<sup>§</sup> | Total (ml) |
| --- | --- | --- | --- | --- |
| 1:16 | 1:7.94 | 17 | 134.98 | 151.98 |
| 2:1 | 1:0.19 | 127 | 24.1 | 151.1 |

<sup>†</sup>Protein:carbohydrate
<sup>‡</sup>Yeast and sucrose solutions were prepared separately and combined in the ratios described to yield each food type.
<sup>§</sup> 200 mL of each solution was prepared with 90 g/L yeast or sucrose, 2 g carrageenan, 1.2 mL propionic acid, and 2 mL Nipagen.

**Supplementary Table S2.** Results of ANOVA on the effect of density (50, 125, 200 larvae per vial), oxygen (15%, 21%, 40% O_2_), nutrition (low and high nutrition diets) and sex on body size (model 1), with all terms and interactions.

| Term<sup>†</sup> | SS | df<sub>Num</sub> | df<sub>Den</sub><sup>‡</sup> | *F*-ratio | *P*-value |
| --- | --- | --- | --- | --- | --- |
| oxygen | 0.011 | 2 | 29.265 | 0.983 | 0.386 |
| nutrition | 4.536 | 1 | 29.408 | 799.206 | **< .001** |
| density | 0.269 | 2 | 29.337 | 23.678 | **< .001** |
| sex | 7.313 | 1 | 2,696.803 | 1,288.469 | **< .001** |
| oxygen:nutrition | 0.045 | 2 | 29.265 | 3.931 | 0.031 |
| oxygen:density | 0.031 | 4 | 29.162 | 1.368 | 0.269 |
| nutrition:density | 0.106 | 2 | 29.337 | 9.364 | **< .001** |
| oxygen:sex | 0.052 | 2 | 2,693.212 | 4.604 | **0.010** |
| nutrition:sex | 0.055 | 1 | 2,696.803 | 9.756 | **0.002** |
| density:sex | 0.013 | 2 | 2,688.960 | 1.172 | 0.310 |
| oxygen:nutrition:density | 0.016 | 4 | 29.162 | 0.691 | 0.604 |
| oxygen:nutrition:sex | 0.007 | 2 | 2,693.212 | 0.575 | 0.563 |
| oxygen:density:sex | 0.051 | 4 | 2,690.068 | 2.243 | 0.062 |
| nutrition:density:sex | 0.000 | 2 | 2,688.960 | 0.039 | 0.961 |
| oxygen:nutrition:density:sex | 0.041 | 4 | 2,690.068 | 1.795 | 0.127 |

<sup>†</sup> Type III sum-of-squares ANOVA.
<sup>‡</sup> Denominator df estimated using Satterthwaite's method.

**Supplementary Table S3.** Results of ANOVA on the effect of density (50 v. 125 larvae per vial), oxygen (21% v. 40%), sex, dataset and survival rate on body size in larvae reared on low nutrition diet.

| Term<sup>†</sup> | SS | df (Num) | df (Den)<sup>‡</sup> | F-ratio | P-value |
| --- | --- | --- | --- | --- | --- |
| oxygen | 0.025 | 1 | 46.413 | 2.378 | 0.130 |
| density | 0.554 | 1 | 43.867 | 52.090 | **< .001** |
| sex | 3.943 | 1 | 1,241.553 | 370.635 | **< .001** |
| survival_rate | 0.059 | 1 | 54.900 | 5.554 | **0.022** |
| dataset | 0.004 | 1 | 39.553 | 0.331 | 0.568 |
| oxygen:density | 0.050 | 1 | 42.154 | 4.692 | **0.036** |
| oxygen:sex | 0.123 | 1 | 1,242.072 | 11.564 | **< .001** |
| density:sex | 0.010 | 1 | 1,241.819 | 0.981 | 0.322 |
| oxygen:density:sex | 0.049 | 1 | 1,242.441 | 4.579 | **0.033** |

<sup>†</sup> Type III sum-of-squares ANOVA.
<sup>‡</sup> Denominator df estimated using Satterthwaite's method.

